# Identifying Causative Mechanisms Linking Early-Life Stress to Psycho-Cardio-Metabolic Multi-Morbidity: The EarlyCause Project

**DOI:** 10.1101/2020.07.08.181958

**Authors:** Nicole Mariani, Alessandra Borsini, Charlotte A.M. Cecil, Janine F. Felix, Sylvain Sebert, Annamaria Cattaneo, Esther Walton, Yuri Milaneschi, Guy Cochrane, Clara Amid, Jeena Rajan, Juliette Giacobbe, Yolanda Sanz, Ana Agustí, Tania Sorg, Yann Herault, Jouko Miettunen, Priyanka Parmar, Nadia Cattane, Vincent Jaddoe, Jyrki Lötjönen, Carme Buisan, Miguel A. González Ballester, Gemma Piella, Josep L. Gelpi, Femke Lamers, Brenda WJH Penninx, Henning Tiemeier, Malte von Tottleben, Rainer Thiel, Katharina F. Heil, Marjo-Riitta Järvelin, Carmine Pariante, Isabelle M. Mansuy, Karim Lekadir

**Affiliations:** Stress, Psychiatry and Immunology Laboratory, Department of Psychological Medicine, Institute of Psychiatry, Psychology & Neuroscience, King’s College London, UK; Department of Epidemiology, Erasmus MC, University Medical Center Rotterdam, Rotterdam, the Netherlands; Department of Child and Adolescent Psychiatry, Erasmus MC, University Medical Center Rotterdam, Rotterdam, the Netherlands; Generation R Study Group, Erasmus MC, University Medical Center Rotterdam, Rotterdam, the Netherlands; Department of Pediatrics, Erasmus MC, University Medical Center Rotterdam, Rotterdam, the Netherlands; Center for Life Course Health Research, Faculty of Medicine, University of Oulu, Oulu, Finland; Medical Research Council Integrative Epidemiology Unit, Bristol Medical School, University of Bristol, UK; Department of Metabolism, Digestion and Reproduction, Genomic Medicine, Faculty of Medicine, Imperial College London, London, UK; IRCCS Istituto Centro San Giovanni di Dio Fatebenefratelli, Biological Psychiatry Laboratory, Brescia, Italy; Department of Psychology, University of Bath, UK; Department of Psychiatry, Amsterdam UMC/Vrije Universiteit & GGZinGeest, Amsterdam Public Health and Amsterdam Neuroscience research institutes, Amsterdam, The Netherlands; European Molecular Biology Laboratory, European Bioinformatics Institute, Wellcome Genome Campus, Hinxton, Cambridge CB10 1SD, United Kingdom (UK); Department of Viroscience, Erasmus Medical Center, Rotterdam, Netherlands; Microbial Ecology, Nutrition and Health Research Group, Institute of Agrochemistry and Food Technology, National Research Council (IATA-CSIC), Valencia, Spain; Université de Strasbourg, CNRS, INSERM, Centre Européen de Recherche en Biologie et Médicine, Institut de Génétique et de Biologie Moléculaire et Cellulaire, PHENOMIN-ICS, Strasbourg, France; Medical Research Center Oulu, Oulu University Hospital and University of Oulu, Oulu, Finland; Department of Information and Communication Technologies, Universitat Pompeu Fabra, Barcelona, Spain; Catalan Institution for Research and Advanced Studies (ICREA), Barcelona, Spain; Universitat de Barcelona, Department of Biochemistry and Molecular Biomedicine, Barcelona, Spain; Department of Social and Behavioral Science, Harvard T.H. Chan School of Public Health, Boston, USA; Empirica Communication and Technology Research, Bonn, Germany; Universitat de Barcelona, Departament de Matemàtiques i Informàtica, Barcelona, Spain; Department of Epidemiology and Biostatistics, MRC-PHE Centre for Environment and Health, School of Public Health, Imperial College London, London, UK; Unit of Primary Health Care, Oulu University Hospital, OYS, Oulu, Finland; Department of Life Sciences, College of Health and Life Sciences, Brunel University London, London, United Kingdom; Laboratory of Neuroepigenetics, Medical Faculty of the University of Zürich and Department of Health Science and Technology of the ETH Zürich, Brain Research Institute, Zürich Neuroscience Center, Switzerland

## Abstract

**Introduction:** Depression, cardiovascular diseases and diabetes are among the major non-communicable diseases, leading to significant disability and mortality worldwide. These diseases may share environmental and genetic determinants associated with multimorbid patterns. Stressful early-life events are among the primary factors associated with the development of mental and physical diseases. However, possible causative mechanisms linking early life stress (ELS) with psycho-cardio-metabolic (PCM) multi-morbidity are not well understood. This prevents a full understanding of causal pathways towards shared risk of these diseases and the development of coordinated preventive and therapeutic interventions.

**Methods and analysis:** This paper describes the study protocol for EarlyCause, a large-scale and inter-disciplinary research project funded by the European Union’s Horizon 2020 research and innovation programme. The project takes advantage of human longitudinal birth cohort data, animal studies and cellular models to test the hypothesis of shared mechanisms and molecular pathways by which ELS shape an individual’s physical and mental health in adulthood. The study will research in detail how ELS converts into biological signals embedded simultaneously or sequentially in the brain, the cardiovascular and metabolic systems. The research will mainly focus on four biological processes including possible alterations of the epigenome, neuroendocrine system, inflammatome, and the gut microbiome. Life course models will integrate the role of modifying factors as sex, socioeconomics, and lifestyle with the goal to better identify groups at risk as well as inform promising strategies to reverse the possible mechanisms and/or reduce the impact of ELS on multi-morbidity development in high-risk individuals. These strategies will help better manage the impact of multi-morbidity on human health and the associated risk.

**Ethics and dissemination:** The study has been approved by the Ethics Board of the European Commission. The results will be published in peer-reviewed academic journals, and disseminated to and communicated with clinicians, patient organisations and media.

## 1. Introduction

### 1.2 Early life stress and psycho-cardio-metabolic multi-morbidity

The World Health Organisation has identified mental disorders, including depression, cardiovascular diseases and diabetes among the six major non-communicable diseases [1]. Individually, each of these groups of diseases represents a burden at the individual and population level. Depression alone is the single largest contributor to global disability in the world, accounting for 12% of total years lived with disability [2] with more than 300 million individuals affected per year. Cardiovascular diseases (CVDs) remain the prime cause of mortality worldwide, accounting for about a third of annual deaths [3]. Finally, type 2 diabetes and related metabolic dysfunctions, including obesity, are a major public health challenge, with an average prevalence of over 8% in the general population [4]. In addition to their separate complexity, existing research has shown important multi-morbidy between these diseases, where multi-morbidity is defined as the co-occurrence of two or more chronic conditions [5]. Epidemiological studies have indeed shown that for example patients experiencing depression are more likely to have comorbid CVD [6], type 2 diabetis [7], or both [8]. However, the specific causative mechanisms leading to psycho-cardio-metabolic (PCM) multi-morbidity are not well understood, which limitslimits the development of effective preventive and therapeutic measures.

Recent evidence suggests that many mental and physical conditions find their origin in the exposure to stress early in life, clinically defined as early-life stress (ELS) [9]. ELS can be both prenatal, such as exposure to clinically-significant depression *in utero*, and postnatal, such as emotional, physical and sexual abuse or neglect in childhood, parental psychopathology and separation, prepubertal bullying, as well as victimisation or violence by peers [10]. Interestingly, growing evidence has supported an association between ELS (both prenatal and postnatal) and the development of the PCM multi-morbidities. Specifically, patients with a history of ELS have higher vulnerability for depression [11], and higher risk of developing cardiovascular disease [12], obesity [13] and type 2 diabetes [14] later in life. Prenatally, the overarching hypothesis is that the maternal stress response is passed to the fetus, via stress hormones crossing the placenta, which affects subsequent brain and physical development of the fetus and newborn [15]. During childhood, exposure to excessive levels of stress early in life can cause several biological alterations which can ultimately favorfavor the development of PCM multi-morbidity [16]. Examples of biological alterations due to response to stress include hypothalamic pituitary adrenal (HPA) axis dysregulation [17], changes in the inflammatory response [18], microbiome dysbiosis [19] and overall bio-psycho-social axis dysfunction [20]. Overall, considering that the prevalence of ELS, both *in utero* and postnatally, whether mild or severe, has reached alarming heights [15], this area of research isessentialis essential for future investigations.

### 1.2 Rationale and overview of the EarlyCause project

EarlyCause is a large-scale inter-disciplinary research that aims to infer evidence for causative mechanisms linking pre- and postnatal ELS to PCM multi-morbidity. It is the product of a collaboration between 14 participating institutions across Europe (Table 1) and is supported by the European Union’s Horizon 2020 research and innovation programme (SC1-BHC-01-2019).

**Table 1.** Participating institutions in the EarlyCause project

| Participant no. | Participant organisation name | Acronym | Country |
| --- | --- | --- | --- |
| 1 (Coordinator) | Universitat de Barcelona | UB | ES |
| 2 | European Molecular Biology Laboratory | EMBL | DE |
| 3 | Erasmus Medical Centre Rotterdam | EMC | NL |
| 4 | University of Zurich | UZH | CH |
| 5 | King's College London | KCL | UK |
| 6 | Consejo Superior de Investigaciones Cientificas | CSIC | ES |
| 7 | Centre Européen de Recherche en Biologie et Médecine | CERBM | FR |
| 8 | University of Oulu | OULU | FI |
| 9 | Fatebenefratelli Institute | IRCCS | IT |
| 10 | University of Bath | UOB | UK |
| 11 | VU Medical Centre, Amsterdam | VUMC | NL |
| 12 | Empirica Communication and Technology Research | EMP | DE |
| 13 | Combinostics Oy | COMBI | FI |
| 14 | Universitat Pompeu Fabra | UPF | ES |

EarlyCause will investigate the hypothesis that ELS, as a risk factor for depressive, cardiovascular and metabolic disorders individually, is a cause of multi-morbidity between these conditions. From a biological point of view, the main hypothesis is that ELS activates a chain of events leading to cellular, molecular, epigenetic and microbial changes which result in dysregulations of processes across tissues. This causative chain would ultimately trigger specific cellular and tissue phenotypes and comorbid pathological traits in the mental, cardiovascular and metabolic domains.

To this end, EarlyCause’s overarching concept is to build upon a unique repertoire of longitudinal data in humans across the lifespan and conduct mechanistic studies in established animal and cellular models to:

i. Identify the causal mechanisms linking exposure to ELS to the risk of multi-morbid symptoms across life course;
ii. Delineate the potential molecular mechanisms underlying these causal associations;
iii. Discover new biomarkers tapping in multiple biological domains;
iv. Build integrative computational models and proof-of-concept tools for multi-morbidity assessment.

The project will focus on four candidate families of biological pathways that have been linked to ELS, specifically:

1. **Epigenetic alterations** are a presumed link between stress exposure and phenotypes. Clear associations between early-life adverse exposure and epigenetic processes (e.g. DNA methylation) and between these epigenetic modifications and later health outcomes have been shown both in humans [21,22, 23]and mouse models [24].
2. **HPA dysregulation** has been identified as a primary biological consequence of ELS exposure [17]. Molecular components of the HPA axis provide a relay chain across the body from the brain to the periphery, and some of the final products (glucocorticoid hormones) are potent regulators of glucose and lipid metabolism. Thus, this represents a central candidate mechanistic player in the aetiology of multi-morbid symptoms.
3. **Inflammatory pathways** are a form of cellular response to ELS [18] reflecting activation of white blood cells in the circulation and peripheral tissues such as the spleen, lymph nodes and adipose tissue. Inflammatory components may have profound effects on the cardiovascular system, endothelial accumulation and activation of plaques, and adipose tissue metabolism, whose dysfunction has been associated with stress-related diseases including depression, cardiovascular disease and diabetes.
4. **Gut microbiome** is a major contributor to health and disease [19] suggested to play a role in modulating immune, neuroendocrine and behavioural responses to ELS, as proven mainly in mouse models [25].

In addition to these biological factors, EarlyCause will test the potential moderating role of key factors such as sex, socioeconomics, and lifestyle in the association between ELS and multi-morbidity development. Evidence for causality, mediation and moderation will be used to identify potential targets for intervention acting on the causative mechanisms to reduce the impact of ELS on multi-morbidity development in high-risk individuals.

## 2. Methods

EarlyCause’s methodology is divided into multiple steps. As shown in *Figure 1*, the study protocol aims to integrate evidence based on (1) longitudinal human data, (2) animal and cellular models, as well as (3) computational bioinformatics and machine learning methods. EarlyCause’s data and methods will be integrated, centralised and standardised, as well as exploited towards the expected impacts.

**Figure 1.**
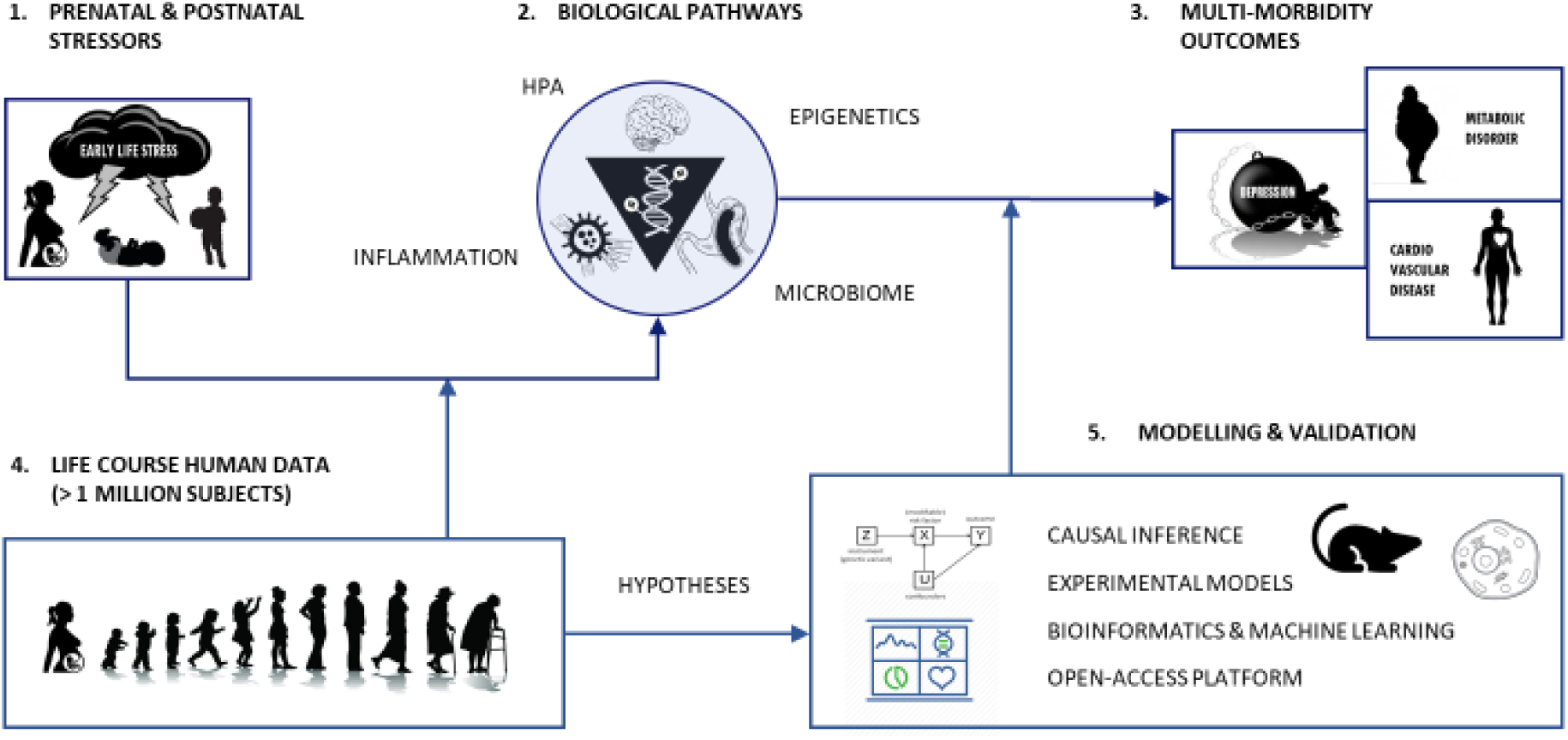
Schematic overview of the EarlyCause project, that will study the link between ELS (1) and multi-morbidity (3), as mediated by biological pathways (2). Large-scale life course human data (4), as well as experimental and computational models will be used to identify and validate the causative mechanisms.

EarlyCause will implement a ‘triangulation’ approach, which will capitalise on the complementary strengths of epidemiological and genetic methods in humans, experimental animal and cellular models, and *in silico* data integration pipelines, as shown in *Figure 2*. This will enable to iteratively and dynamically:

- Apply association analyses and latent modelling to large-scale longitudinal human data on ELS, resulting in the identification of potential new candidate biomarkers of PCM multi-morbidity, as well as novel hypotheses on underlying causative mechanisms;
- Apply causal inference methods, including structural equation modelling, Mendelian randomisation, and molecular mediation, to infer the causal relationships between ELS, biological mediators and the multimorbid outcomes;
- Validate the mechanisms and identify the associated molecular pathways in pre- and postnatal animal models, using established cellular models of stress;
- Integrate the identified determinants and molecular markers of ELS into computational models of multi-morbidity across the life span, andand design a proof-of-concept decision support tool for PCM multi-morbidity risk assessment, by extending an existing single-disease e-health tool commercialised by EarlyCause partner COMBI (*i.e*., from the DSF^®^: *Disease State Fingerprint [26,27]* to the MSF: *Multi-morbidity State Fingerprint*).

**Figure 2.**
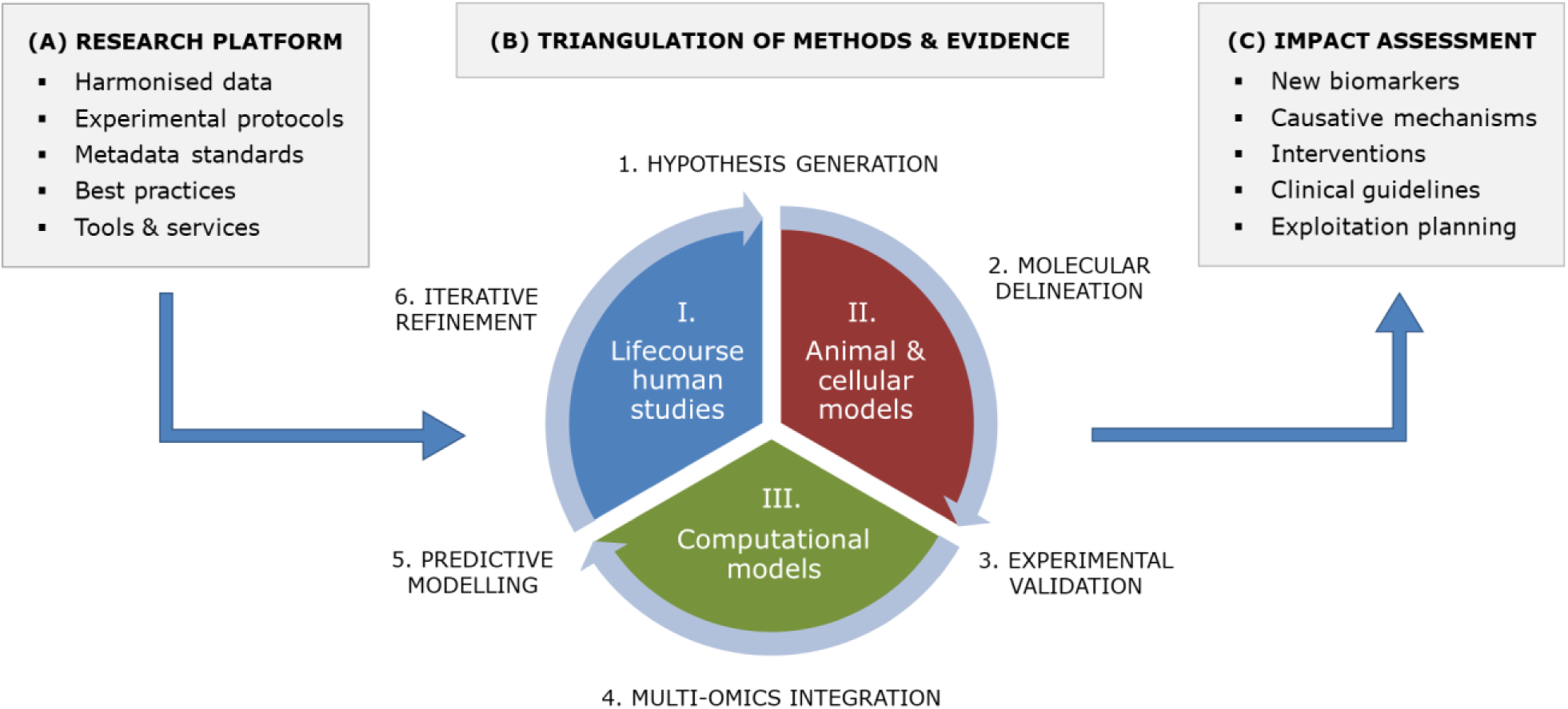
Overall methodology of the EarlyCause project, including (A) research platform to use of harmonised data from existing resources, research protocols and best practices; (B) triangulation approach, which will capitalise on the complementary strengths of epidemiological and genetic methods in humans, experimental animal and cellular models, and in silico data integration pipelines; and (C) expected results such as new biomarkers and clinical knowledge.

### 2.1 Longitudinal human data

#### a. Hypothesis-generating analyses of biological markers & environmental moderators

We will leverage harmonized data from a large set of human studies to examine the relationship between ELS and multi-morbidity across the lifespan, identify potential molecular markers and quantify the protective vs. exacerbating role of modifiable lifestyle factors. These datasets together span from pregnancy to old age, including the well-known Avon Longitudinal Study of Parents and Children (ALSPAC), Generation R Study (GenR), Northern Finland Birth Cohorts (NFBC), UK Biobank, Rotterdam Study, and the Netherlands Study of Depression and Anxiety (NESDA). We will make use of correlational multivariate analyses as well as novel latent modelling techniques to model the shared versus unique contribution of ELS on multi-morbid outcomes (*Figure 3*). In these human studies, we will:

- Define the relationship between ELS and multi-morbidity across the lifespan, by tracking risk factors of cardiovascular and metabolic disorders in children and adolescents, and the influence of early life stressors on tracking patterns, drawing on a rich datasets of clinical samples and cohort studies publicly available or through members and collaborators of the consortium. We are currently seeking additional scientists and groups interested in collaborating with us;
- Identify candidate biological predictors and mediators of ELS effects on multi-morbidity (epigenetic marks, neuroendocrine function, inflammation, gut microbiome);
- Quantify the protective or exacerbating role of modifiable lifestyle factors (e.g. exercise, diet, sleep, smoking, alcohol use) in the relationships of ELS with biological markers and PCM multi-morbidity;
- Provide hypotheses and candidate biomarkers that can be used for causal inference and mediation studies, as well as in animal and *in vitro* studies;

**Figure 3.**
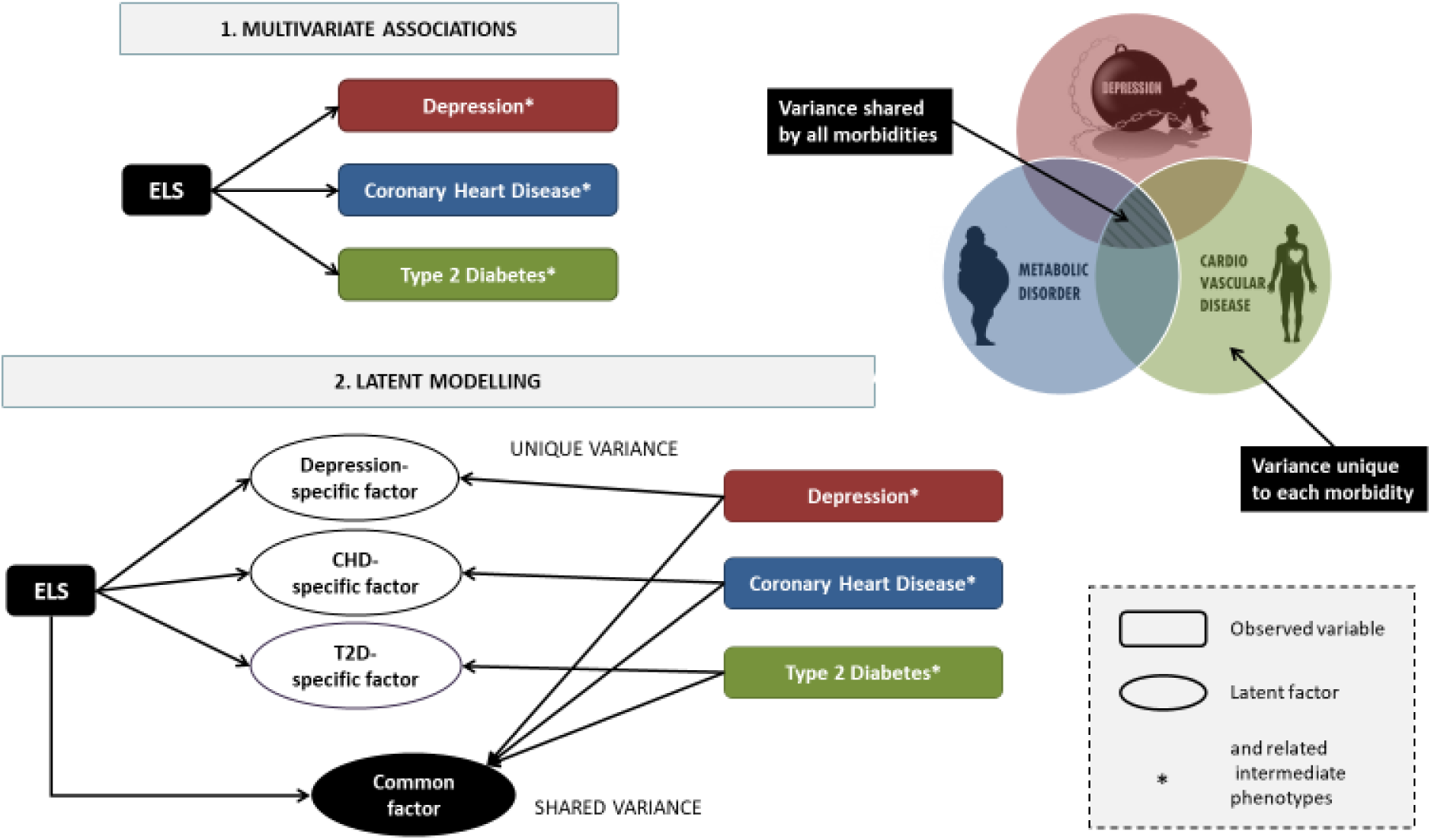
Differences between (1) simple multivariate analysis and (2) latent modelling of psycho-cardio-metabolic multi-morbidity implemented in EarlyCause.

#### b. Causal inference and molecular mediation analyses

To investigate whether ELS represents a *causal* risk factor for PCM multi-morbidity, and to what extent biological factors identified in the hypothesis-generating analyses represent shared non-causal biomarkers of ELS and PCM multi-morbidity, or point towards causal mediating mechanisms, we will use multiple approaches. We will apply multiple methods (triangulation) such as Mendelian randomisation, genetic risk score methods and associated sensitivity analyses [28] to infer causality using population-based human genetic data. Genetic summary measures on ELS and multi-morbidity will be derived through meta-analysis of genome-wide association study (GWAS) data on childhood maltreatment as well as on health outcomes (i.e. depression, type 2 diabetes, and coronary heart disease). More specifically, we will:

- Establish a catalogue of genetic instrumental variables for ELS by performing a GWAS-meta-analysis across studies of human cohorts involved in EarlyCause and, if possible, including further studies with relevant data;
- Infer the causal association between postnatal ELS and multi-morbidity development through Mendelian randomisation and by using both diagnostic criteria and pre-diagnostic correlates of multimorbid outcomes;
- Establish the molecular mediation of biological markers (DNA methylation, cortisol, inflammation, microbiome) linking ELS exposure to later PCM multi-morbidity.

### 2.2 Animal and cellular models

#### a. Modelling ELS in animals for causality assessment

We aim to exploit unique pre- and postnatal rodent models of stress [24,29] to identify ELS-associated molecular pathways causally linked to multi-morbid symptoms in adult life. For both models, we will determine which changes in the epigenome, transcriptome and proteome/metabolome are induced by ELS exposure across different tissues and biological fluids relevant for PCM symptoms, including brain, blood, heart, liver, pancreas and adipose tissue (*Figure 4*). The purpose will be to identify common and distinct epigenetic and neuroendocrine factors, immune markers and molecular pathways dysregulated by ELS in these tissues in the animal models. The observed alterations will be cross-validated with markers/pathways identified in humans via comparative analyses. Once epigenetic, neuroendocrine, immune and molecular alterations are identified, their potential reversibility by interventions such as environmental enrichment or pharmacological compounds will be assessed, based on previous knowledge that enriched life conditions have beneficial effects on brain and body functions. In avalidation step, we will examine the causal involvement of relevant markers in the aetiology and expression of symptoms characteristic of depression, cardiovascular dysfunctions and metabolic dysregulation by experimental manipulations *in vivo*. A final aspect will be to examine the gut microbiome composition in association with ELS exposure in the animal models and the relationships with other molecular alterations, and compare the findings with those in humans. The implication of the human gut microbiome in the development of multi-morbidity symptoms will also be tested by microbiota transplantation experiments into rodents and phenotypic analyses. Specifically, we will:

- Determine and quantify the impact of pre- and postnatal stress on behavioural, cardiovascular and metabolic functions in adulthood in rodents;
- Examine the effects of intervention and identify moderators relevant for humans;
- Identify epigenetic and molecular pathways associated with symptoms, and test causality *in vivo*;
- Assess the specific role of the human gut microbiome as causative factor for the development of PCM multi-morbidity.

**Figure 4.**
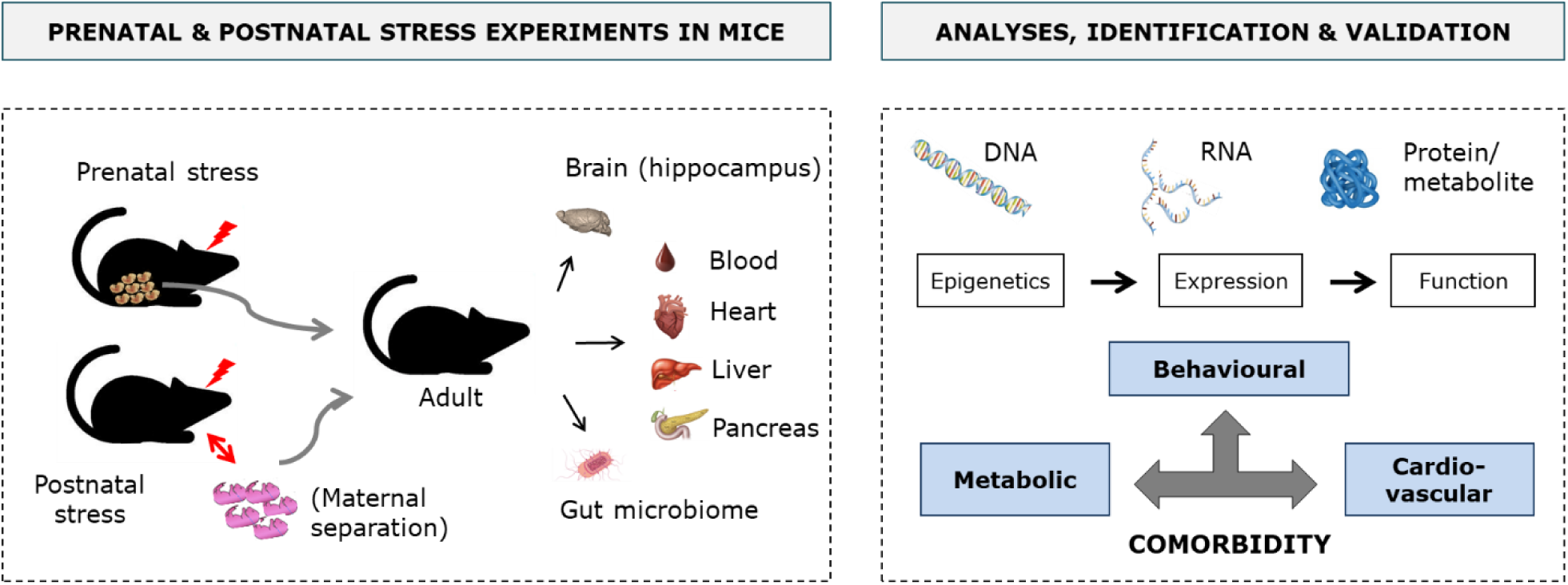
Overview of the experiments and analyses in rodents to identify molecular pathways linking ELS to PCM multi-morbidity.

#### b. Cellular models to identify causal molecular mechanisms of ELS-induced multi-morbidity

We aim to uncover causative molecular and biological mechanisms underpinning ELS-associated multi-morbidity between depression, coronary heart disease and diabetes type 2 associated with ELS by leveraging a variety of human cellular models (*Figure 5*). In particular, we will use a coherent, systematic approach to mimic ELS-relevant insults across cells from the human brain, heart, liver, pancreas, and blood immune system, to identify the molecular processes induced by ELS that influence cellular and tissue homeostasis, resulting in multi-morbid symptoms. Specifically, we aim to:

- Establish *in vitro* conditions to mimic stress and metabolic insults in human cell lines and primary cultures derived from brain, heart, liver, pancreas, and blood immune system;
- Study the effects of ELS on different cellular phenotypes related to depression, coronary heart disease, and diabetes type 2;
- Test causal mechanisms based on candidates’ biomarkers obtained from human and animal studies through molecular manipulations in selected cellular systems;
- Develop novel research tools and experimental strategies to study the underlying mechanisms in the communication between blood and other tissues after ELS, and to model sex effects;
- Identify the molecular signature of ELS in the distinct cellular types.

**Figure 5.**
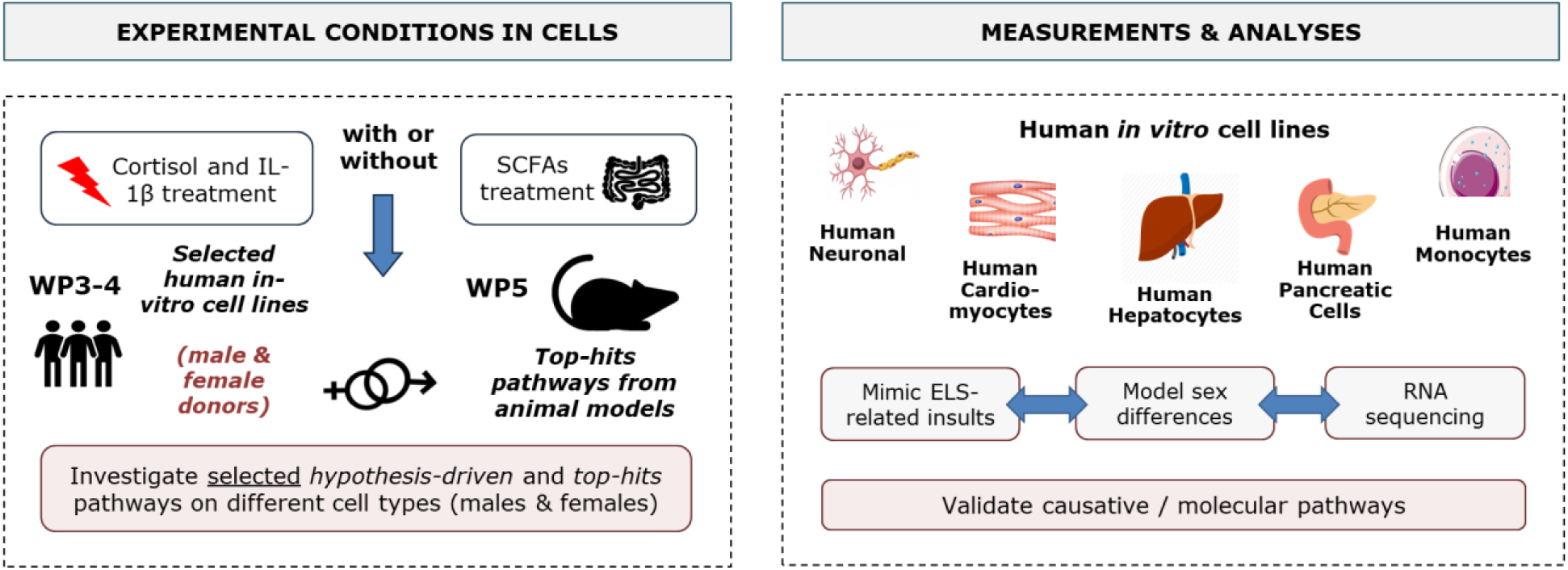
Overview of the cellular modelling experiences. SCFAs (Short-chain fatty acids)

### 2.3 Computational bioinformatics and machine learning methods

We plan to implement advanced bioinformatic, statistical and machine learning techniques to integrate and leverage the findings and determinants derived from human studies and experimental models. Several types of integration will take place:

- Multi-omics integration of molecular interaction networks at different levels (DNA, RNA and proteins/metabolites) to dissect out the mechanistic chains across tissues;
- Structural equation modelling to model developmental timing and direction of associations, i.e. direct effects, as well as indirect pathways between variables and lifestyle factors affecting the pathways;
- Multi-cohort integration for bridging child/adolescent, adult and elderly cohorts and thus offer a life-course perspective on the link between ELS and multi-morbidity development;
- Machine learning models of multi-morbidity using unsupervised deep learning to simulate patient-specific trajectories towards multi-morbidity integrating identified biomarkers and pathways.

Subsequently, a proof-of-concept software will be assembled by integrating the predictive models within an existing e-health tool commercialised by COMBI; the Disease-State Fingerprint (DSF®). EarlyCause will extend it to account for multi-disease data and associations for the first time. The obtained tool will be pilot tested by COMBI’s usability experts to assess its acceptance and potential in future clinical management of multi-morbidity.

### 2.4 Centralised research platform

The research proposed in EarlyCause is novel, integrating causal inference studies, experimental models of both pre- and postnatal stress, and new computational approaches for uncovering the causal effects of ELS on multi-morbidity development. The expected results, including those on the role of epigenetics, microbiome and environmental modifiers, will set the stage for new studies to generate knowledge and contribute to public health guidelines. We aim to establish a research-enabling web-platform that will integrate data services, experimental standards and best practices to support next-generation research on ELS and multi-morbidity. The EarlyCause web-portal and centralised platform represented in *Figure 6* will provide a comprehensive support to researchers, which will allow them to upload, search and ecurate data relating to ELS-induced multi-morbidity. For full FAIR (Findable, Accessible, Interoperable and Re-usable) compliance, our strategy is to build upon existing life-science/data infrastructures such as ELIXIR [30] and the EMBL European Bioinformatics Institute.

A key feature of the web-portal will be a rich environment for the discovery and selection of appropriate data sets and relevant project protocols for further exploration. These functions will leverage and adapt the existing data hub/portal software framework as developed and used in such projects as COMPARE [31] and HipScI [32]. The EarlyCause web-portal will be linked to ELIXIR’s “core” and “deposition” databases, notably the European Nucleotide Archive [33] (ENA) and European Genome-phenome Archive [34] (EGA) for fully open and controlled access molecular data, respectively, as well as BioSamples [35] for sample-related data, such as ELS exposure, rodent model stress descriptors, and Biostudies [36] for a variety of assay data types, such as rodents behavioural data and metabolic profiling. For image data, in particular rodents histology, we will leverage the image database from euro-BioImaging [37].

**Figure 6.**
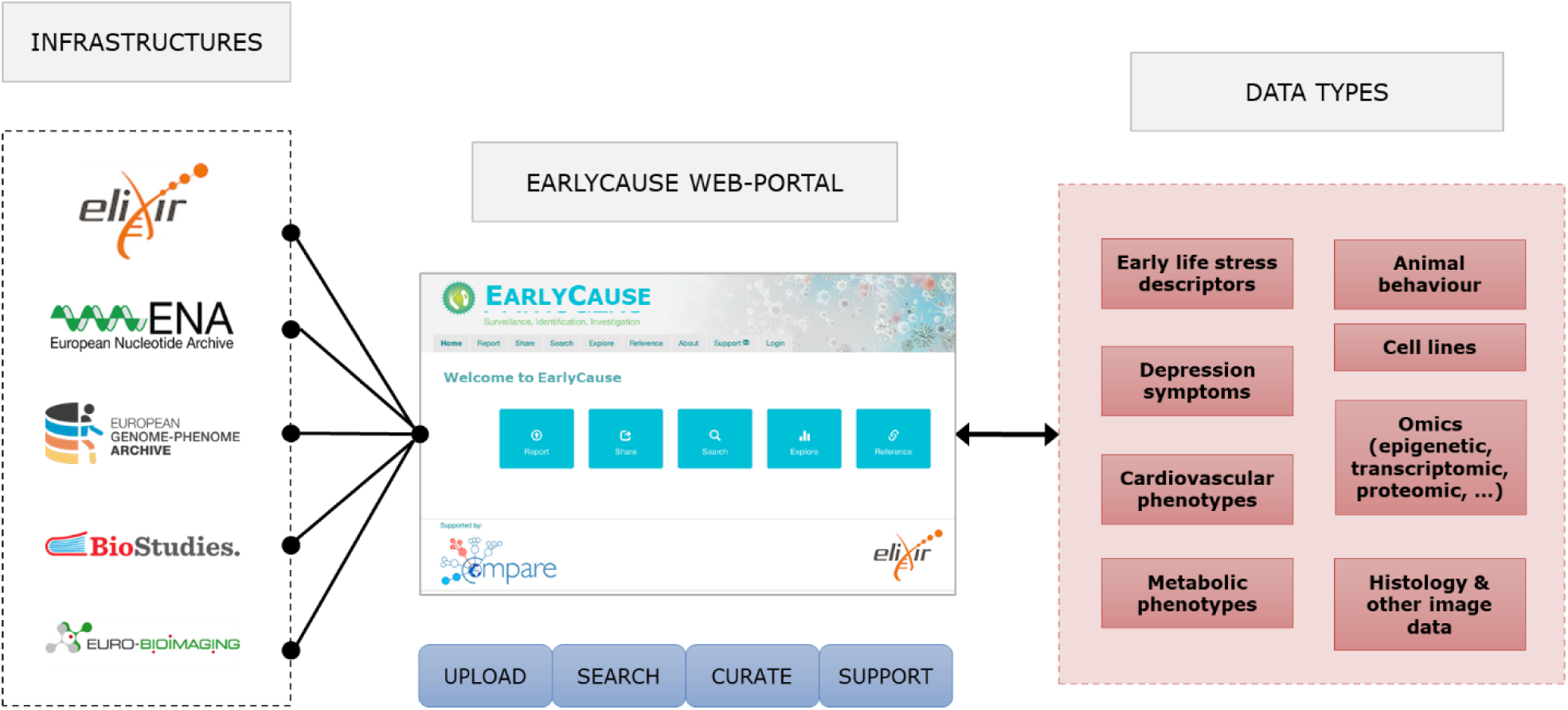
Overview of the EarlyCause centralised research platform, which will allow to upload, search and manage human and experimental data for investigating ELS-induced multi-morbidity.

### 2.5 Impact assessment and exploitation planning

Since the study of ELS and its effects on multi-morbidity represents a novel research field, the EarlyCause consortium will perform a thorough impact evaluation, spanning socioeconomics, healthcare practice, prevention strategies, as well as technology and market analysis. The analysis will be built upon the ASSIST tool-kit, which will use quantitative input from literature- and expert-informed data, established socioeconomic models and Monte-Carlo simulation, to perform a qualitative analysis for different stakeholders, e.g. users, beneficiaries, payers, technologists, organisations or health-systems. The experience obtained in impact assessment of the C3-Cloud EU project [38], which developed clinical decision supports for the management of multimorbid chronic patients, will strengthen these activities. For healthcare practice, ex-ante scenarios will be designed for three countries (Germany, Spain and the UK) and compared to as-is situations to assess potential impact of the research findings (new biomarkers, causal mechanisms, specific role of modifiers such as microbiome) on multi-morbidity screening and prevention. The resulting evidence-based impact assessment will contribute to the accelerated diffusion of project results and their acceptance by the social care, healthcare, and policy communities and facilitate future research activities.

## 3. Discussion

Overall, EarlyCause will explore new territories at the interface of fundamental and clinical research by addressing the question of how ELS biologically impacts PCM multi-morbidity development. This will provide a rich series of translational research lines for targeting prevention, diagnosis, prognosis, therapy development, and management of PCM multi-morbidity (*Figure 7*).

### New directions for prevention and diagnosis of ELS-induced multi-morbidity

EarlyCause aims to create knowledge about the causal impact of ELS on multi-morbidity with the goal to inform the development of prevention programmes in two main directions. The first direction concerns the allocation of resources to schemes focused on reducing ELS per se, such as by providing greater support to high-risk families during pregnancy (e.g. midwife support, family and school-based programmes), or by increasing resilience to ELS through supporting early emotional, behavioural, and physical regulation in children. The second research direction concerns the identification of relevant targets for preventing multi-morbidity itself. Information on (the direction of) causality will allow the most effective primary preventative strategies to be established. This might focus on promoting lifestyle changes that affect possible shared causes of multi-morbidity, or treating the primary cause directly, or preventing/treating all multimorbid conditions together. In addition, knowledge of the role of ELS in PCM multi-morbidity development can also enhance the identification of multimorbid conditions in patients screened as having been exposed to ELS and who have already been diagnosed with one disorder (e.g. depression, but not yet diabetes or coronary heart disease). Furthermore, EarlyCause will combine ongoing research lines in a unique framework between ELS, inflammation, HPA, and microbiome, which will be scaled-up and extended to include different ‘omics’ levels (e.g. microbiome, genomics, epigenetics). This will open new routes to diagnose multi-morbidity beyond the simple addition of traditional symptom-based categories, promoting the development of a more biologically-informed nosology of multi-morbidity.

### Therapy development

EarlyCause will also promote new research for identifying targets for intervention. A natural next step will establish whether known drugs can impact the identified biomolecular pathways (so-called drug repurposing). This will open a host of potential future clinical trials using repurposed drugs that target these specific mechanisms. Randomised controlled trials are the gold standard for obtaining evidence on the effects of modifying disease risk processes. However, traditional drug (repurposing) development has several limitations, including short follow-up, small sample size, and non-representative samples. In this case, our Mendelian randomisation-based findings on PCM multi-morbidity can have direct implications for drug repurposing or the identification of unintended drug side effects.

### Management of multi-morbidity

Finally, knowledge gathered from EarlyCause will open opportunities for developing new patient pathways and care models for addressing ELS-related PCM multi-morbidity, complemented with an innovative set of technical solutions for improved clinical decision-making. The key aim of EarlyCause is also to identify lifestyle factors that dampen or exacerbate the impact of ELS on PCM multi-morbidity risk. Such knowledge will impact the implementation of lifestyle changes that can ameliorate symptoms and disease course, particularly amongst those who have already been exposed to ELS. EarlyCause will therefore improve existing clinical guideline recommendations with economic modelling of benefit and harm. Our ideal end-point will be to publish generated evidence to inform the future development of more streamlined and optimised multi-morbidity care pathways, thus improving decision-making and clinical management of patients with ELS-related PCM multi-morbidity.

**Figure 7.**
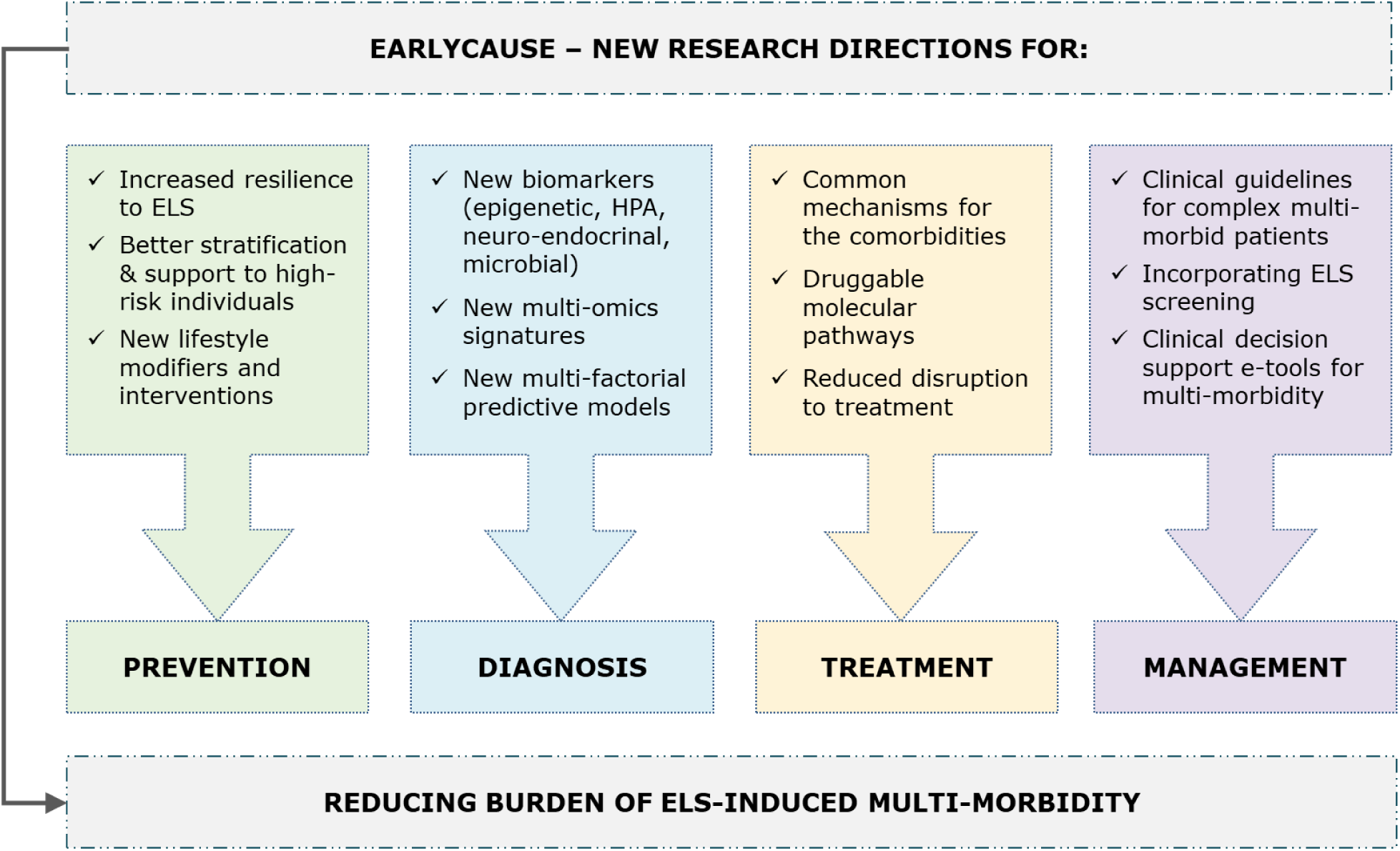
Overview of the research directions affected by EarlyCause.

## 4. Conclusion

In the next years, EarlyCause will establish extensive research linking human, animal and cell studies with the aim to clarify how ELS biologically impacts PCM multi-morbidity development. The consortium will operate on FAIR data management and open science practice aiming to impact on diagnostic tools and new health policies to alert on ELS and prevent its life long consequences.

## 5. Acknowledgments

This work is supported by the European Union’s Horizon 2020 research and innovation programme (grant n° 848158). All the authors have contributed equally to this paper.

## Notes

### Competing Interest Statement

The authors have declared no competing interest.

